# Myt1 kinase promotes mitotic cell cycle exit in *Drosophila* intestinal progenitor cells

**DOI:** 10.1101/785949

**Authors:** Reegan J. Willms, Jie Zeng, Shelagh D. Campbell

**Affiliations:** Department of Biological Sciences, University of Alberta, Edmonton, Alberta T6G 2E9, Canada

## Abstract

Inhibitory phosphorylation of Cdk1 is a well-established mechanism for gating mitotic entry during development. However, failure to inhibit Cdk1 in adult organs causes ectopic cell division and tissue dysplasia, indicating that Cdk1 inhibition is also required for cell cycle exit. Two types of progenitor cells populate the adult *Drosophila* midgut: intestinal stem cells (ISCs) and post-mitotic enteroblasts (EBs). ISCs are the only mitotic cells under homeostatic conditions, dividing asymmetrically to produce quiescent EB daughter cells. We show here that Myt1, the membrane associated Cdk1 inhibitory kinase, is required for EB quiescence and subsequent differentiation. Loss of Myt1 disrupts EB cell cycle dynamics, promoting Cyclin A-dependent mitosis and accumulation of smaller progenitor-like cells that fail to differentiate. Thus, Myt1 inhibition of Cyclin A/Cdk1 functions as a mechanism for coupling cell cycle arrest with terminal cell differentiation in this developmental context.

## INTRODUCTION

An equilibrium between cell division, differentiation and loss is essential for the establishment and maintenance of tissue homeostasis in adult animals. Most tissues possess progenitor cell populations that proliferate to produce specialized cell types. These progenitors rely upon Cdk (<u>c</u>yclin-<u>d</u>ependent <u>k</u>inase) activity to drive the mitotic cell cycle, a process that must be tightly controlled to promote tissue renewal yet avoid over-proliferation. As progenitors exit the mitotic cell cycle and begin to differentiate, Cdk1 is inhibited by several mechanisms. These include expression of Cyclin dependent kinase inhibitors (CKIs) and APC/C mediated degradation of mitotic cyclins (Buttitta and Edgar, 2007; Pimentel and Venkatesh, 2005). Some differentiating cells also inhibit Cdc25 phosphatase, which promotes mitosis by removing Cdk1 phosphorylation catalyzed by Wee1-like kinases (Schaeffer et al., 2004; Shcherbata et al., 2004). However, a direct mechanism linking Cdk1 inhibitory kinase activity to mitotic cell cycle exit has not yet been described.

Two Cdk1 inhibitory kinases exist in *Drosophila,* Wee1 and Myt1. Previous genetic studies have defined specialized developmental functions for each kinase. Maternally-provided Wee1 activity is essential for a DNA replication checkpoint that prevents mitotic catastrophe during the rapid cleavage cycles of early embryogenesis (Fasulo et al., 2012; Price et al., 2000; Stumpff et al., 2004). Myt1 kinase has a broader range of functions that are essential at multiple stages of development. These include male pre-meiotic G2 phase arrest (Varadarajan et al., 2016), pre-mitotic checkpoint responses to DNA damage in imaginal discs (Jin et al., 2008), and mitotic cell cycle exit of germline-associated somatic cells (Jin et al., 2005). The exact mechanism by which Myt1-mediated Cdk1 inhibitory phosphorylation promotes quiescence in differentiated cell types remains unclear.

To address this issue, we have genetically characterized Cdk1 inhibitory phosphorylation in progenitor cells of the *Drosophila* intestinal epithelium. In this system, self-renewing stem cell progenitors (ISCs) divide asymmetrically to produce enteroblasts (EBs), a transient post-mitotic cell type that differentiates into absorptive enterocytes (ECs) (Micchelli and Perrimon, 2006; Ohlstein and Spradling, 2006; Ohlstein and Spradling, 2007). We found that Myt1 inhibitory phosphorylation of Cyclin A/Cdk1 is required to block proliferation of normally post-mitotic EBs. Furthermore, Myt1 loss prevents proper EC specification, demonstrating that Cdk1 inhibitory phosphorylation is required for coordinating mitotic cell cycle exit with cell differentiation.

## RESULTS

### Myt1 regulates cell division of intestinal progenitors

Compromising Cdk1 inhibitory phosphorylation causes ectopic cell proliferation during *Drosophila* development, notably by loss of Myt1 function (Ayeni et al., 2016; Jin et al., 2005; Jin et al., 2008). To investigate the hyperproliferative phenotype of *myt1* mutants, we assessed its role in the adult fly intestine, a tissue where mechanisms of progenitor division and differentiation have been well characterized. Myt1 is one of two distinct Cdk1 inhibitory kinases, so we initially analyzed cell proliferation in both *wee1* (Price et al., 2000) and *myt1* (Jin et al., 2005) null mutants, using immunolabeling of phospho-histone H3 (PH3) to mark mitotic cells (Hendzel et al., 1997). Proliferating cells were rarely observed in control intestinal epithelia, as expected under homeostatic conditions (Figures 1A and 1D). In contrast, there were roughly 5-fold more mitotic cells in *wee1* mutants (Figures 1B and 1D) and 40-fold more mitotic cells in *myt1* mutants (Figures 1C and 1D). Hyper-proliferation in the *myt1* mutants occurred as early as the 3^rd^ instar larval stage (Figures S1A and S1B). Although both Wee1 and Myt1 clearly influence proliferation, the extreme defects observed in *myt1* mutants implicates Myt1 as the predominant Cdk1 inhibitory kinase with respect to epithelial homeostasis.

**Figure 1.**
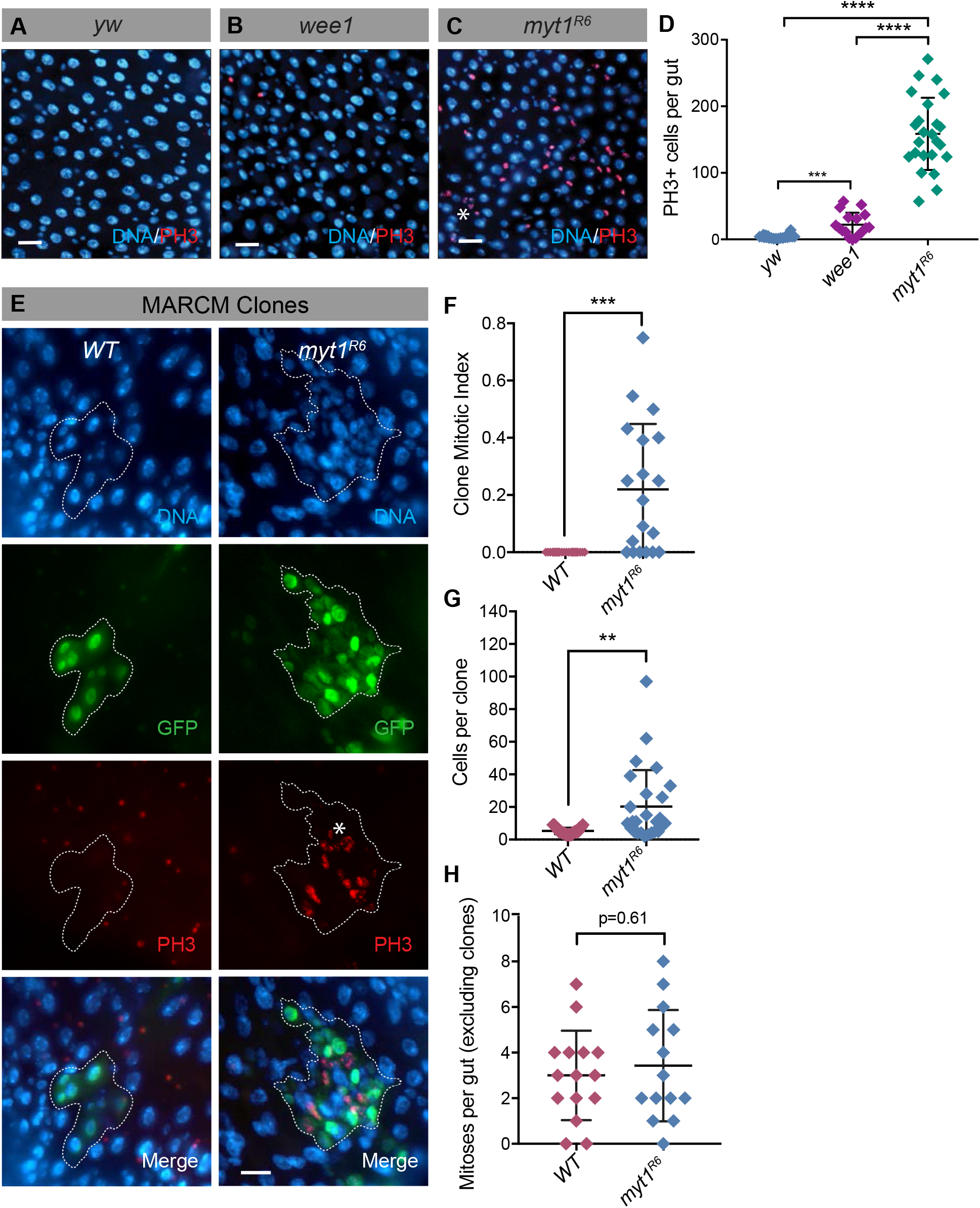
Increased proliferation in the intestinal epithelium of *myt1* mutants. **A)** Region of *yw* posterior midgut showing no observable mitoses. Asterix indicates PH3-positive puncta. Scale bar = 20 µm. **B)** Several mitotic cells are observable in this section of *wee1* mutant posterior midgut. Scale bar = 20 µm. **C)** Section of *myt1* mutant posterior midgut showing PH3-positive cells. Scale bar = 20 µm. **D)** Quantification of PH3-positive cells (mitoses) per whole gut. Genotypes: *yw* (n=22), *wee1* (n=17) and *myt1* (n=24). (t test: *** p<0.0005, **** p<0.00005). **E)** Posterior midgut with WT or *myt1* MARCM clone (GFP-positive) outlined. Asterix indicates PH3-positive puncta. Scale bar = 20 µm. **F)** Comparison of mitotic indices in WT (n=24) and *myt1* mutant (n=19) clones. (t test, *** p<0.0005). **G)** Cells per clone in WT (n=27) and *myt1* MARCM clones (n=26). (t test, ** p<0.005). **H)** Mitoses per gut (WT, n=15; *myt1/+,* n=14) excluding mitoses within WT or *myt1* clones.

To determine if Myt1 was operating cell autonomously we performed genetic mosaic analysis, using the repressible cell marker (MARCM) system to generate GFP-positive *myt1* mutant clones from ISCs (Lee and Luo, 1999). In wild-type controls, mitotic cells were never observed at 7 days after clone induction (Figures 1E and 1F) and there was little variation in the number of cells per clone (Figure 1G). In contrast, mitotic cells were often observed in *myt1* mutant clones (Figures 1E and 1F) and these clones contained many smaller cells relative to controls (Figure 1E and 1G). We also noted that PH3-labeling of nuclei appeared punctate in *myt1* mutants, rather than uniform as in controls (Figures 1C and 1E). Similar mitotic chromatin defects were previously described in *myt1* mutant imaginal wing discs (Jin et al., 2008). Closer examination of fixed mitotic cells revealed an apparent increase in the numbers of cells in prophase relative to controls, however cells at all stages of mitosis were present (Figures S1C and S1D). This result, along with the increased numbers of cells in *myt1* mutant clones, demonstrates that *myt1* mutant cells are able to complete cell division. There was no effect on tissue proliferation surrounding the *myt1* clones (Figure 1H). Myt1 therefore operates cell-autonomously to regulate cell proliferation and ensure homeostasis in the adult intestinal epithelium.

### Myt1 promotes mitotic cell cycle exit of enteroblasts

Differentiated epithelial cells and visceral muscle (VM) provide mitogenic cues to ISCs (Jiang et al., 2009; Lee et al., 2009; Lin et al., 2008). Although our clonal analysis showed that Myt1 functions cell-autonomously, it remained unclear whether Myt1 intrinsically regulates progenitor proliferation or transduces signals from a differentiated cell type. To address this issue, we examined which cell type(s) were affected by Myt1 depletion using transgenic, temperature-dependent expression of *myt1^RNAi^*. To empirically determine if RNA interference was effective for this experiment, we expressed EGFP-Myt1 (Varadarajan et al., 2016) with or without *myt1^RNAi^*, using *esgGAL4^ts^* to drive transgene expression in both ISCs and EBs (Buchon et al., 2009). Several *myt1^RNAi^* lines were initially tested that gave similar results and one line was chosen for subsequent analysis (VDRC 105157). The EGFP-Myt1 signal localized primarily to the nuclear envelope of control progenitor cells after 7 days transgene induction, whereas *myt1^RNAi^* co-expression eliminated this signal (Figure S2A). Myt1 protein levels were therefore effectively depleted by this protocol. When *myt1^RNAi^* was expressed independently in either ISCs or EBs, there were significantly more mitotic cells in the posterior midgut relative to controls (Figure 2A). Expression of *myt1^RNAi^* in enteroendocrine cells (EEs), ECs, or VM had no effect, however (Figure 2A). Myt1 activity is therefore required specifically in ISCs and EBs for normal proliferation kinetics.

**Figure 2.**
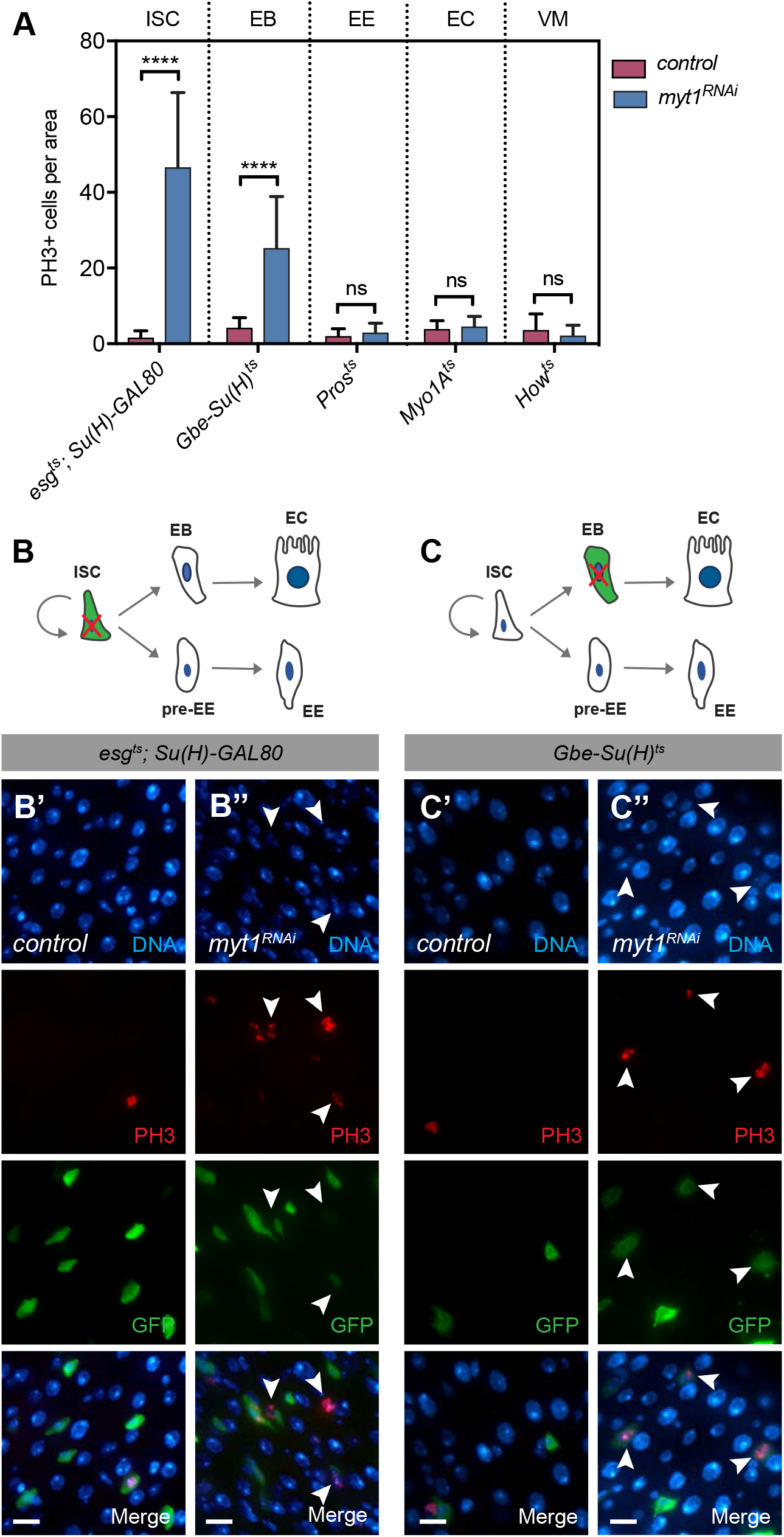
Myt1 loss drives EBs into mitosis. **A)** Temperature-sensitive GAL4 drivers specific to each epithelial cell type were used to analyze Myt1 requirements in flies that were kept at 29°C for 7 days before dissection. *esg^ts^; Su(H)-GAL80* (ISC-specific) and *GBE^ts^* (EB-specific) depletion of Myt1 resulted in significant increases in mitoses (n=21 and 24 guts, respectively) relative to expression of UAS-GFP alone (n=16 and 16 respectively). *Pros^ts^*, *Myo1A^t^*^s^, and *How^ts^* driving *myt1^RNAi^* (n=18, 25, and 19 respectively) in EE, EC and VM showed no significant change in mitoses relative to expression of UAS-GFP alone (n=17, 19, and 16 respectively). Analysis was done on projected Z-stack images of guts taken 1 frame anterior from the midgut-hindgut transition zone. (t test, **** p<0.00005). Error bars represent +/-SD. **B-C)** Visualization of mitosis upon Myt1 depletion in progenitor cells. Schematics depict cells depleted of Myt1 with a red X, while cells expressing GFP are in green. **(B)** Control shows an example of mitosis in a GFP-positive cell (**B’**), while Myt1-depleted ISCs have PH3 present in GFP-negative cells **(B”). (C)** Mitosis is never observed in GFP-positive EB controls **(C’)**, however EB-specific Myt1 depletion results in mitotic EBs (**C”**, arrowheads). Scale bars = 10 µm.

Next, we more closely examined ISCs and EBs that were independently depleted of Myt1. We used ISC-specific or EB-specific GAL4 lines driving a GFP reporter to identify transgene-expressing cell types (Wang et al., 2014; Zeng et al., 2010). In control epithelia expressing the ISC reporter alone (Figure 2B), all mitotic cells were clearly GFP-positive and therefore identifiable as ISCs (Figure 2B’). In contrast, mitotic cells co-expressing ISC-specific *myt1^RNAi^* were often weakly GFP-positive or GFP negative (Figure 2B”). This could mean that Myt1 loss reduced GFP reporter expression or that it promoted mitotic activity of a secondary cell type. To explore the latter possibility, we tested whether ISC-specific Myt1 depletion promotes ectopic EB mitosis. For this experiment we co-expressed an EB reporter consisting of the Notch response element (NRE) tagged with GFP (Lucchetta and Ohlstein, 2017; Ohlstein and Spradling, 2007) with ISC-specific *myt1^RNAi^*. Almost half of the mitotic cells observed were NRE-positive (Figures S2B and S2C), showing that ISC-specific Myt1 depletion produced mitotic EBs. We then analyzed EB-specific expression of *myt1^RNAi^* using the *GBE-Su(H)^ts^* driver. Mitotic EBs were never observed in controls expressing GFP alone (Figure 2C’), whereas expression of *myt1^RNAi^* produced EBs that were both GFP and PH3-positive (Figure 2C”). We therefore conclude that Myt1 activity is required in both progenitor cell types to prevent ectopic mitosis of naturally quiescent EBs.

### Myt1 loss alters enteroblast cell cycle kinetics

*Drosophila* Myt1 has previously been characterized as a regulator of G2 phase arrested cells (Jin et al., 2008; Varadarajan et al., 2016). We therefore suspected that an inability to arrest in G2 phase might be responsible for ectopic proliferation of Myt1-depleted EBs. To test this idea, we examined how loss of Myt1 activity affected cell cycle timing using the fly-FUCCI system that differentially labels cells at different stages (Zielke et al., 2014). Consistent with previous data (Kohlmaier et al., 2015; Zielke et al., 2014), roughly 43%, 2%, and 48%, of control progenitor cells were in G1, S, and G2/M phases respectively, with 10% being unclassifiable due to weak fluorescent signal intensity (Figures 3A and 3C). RNAi depletion of Myt1 in both ISCs and EBs reduced the number of G1 phase cells to 21% and virtually eliminated S and G2/M phase reporter labeling, with most cells being unclassifiable (Figures 3B and 3C). Notably, the Cyclin B reporter was absent from Myt1-depleted progenitors (Figure 3B), which is consistent with these cells failing to arrest properly in G2 phase when Cyclin B normally accumulates (Lehner and O’Farrell, 1990).

**Figure 3.**
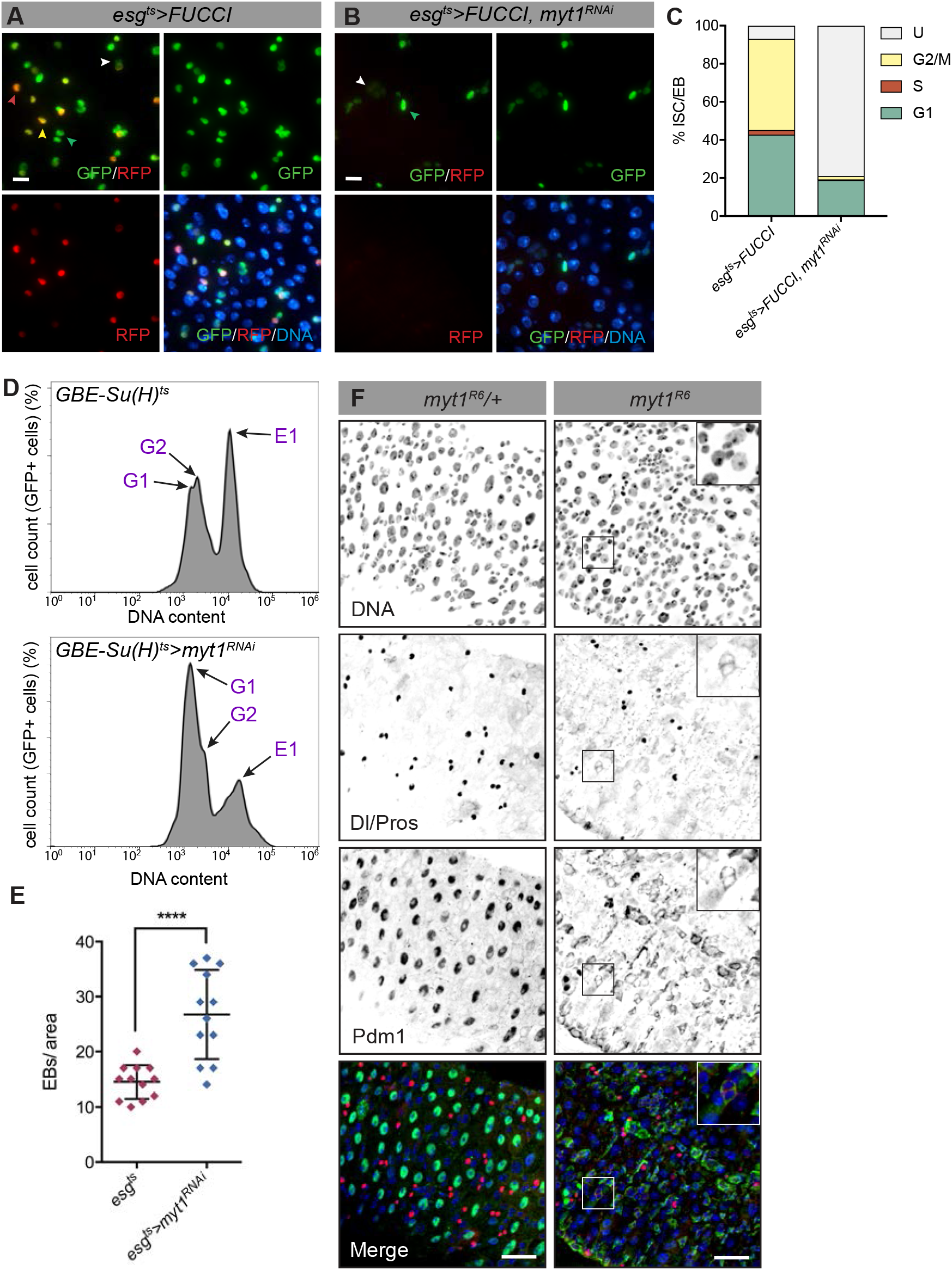
Cell cycle and cell differentiation are disrupted in Myt1-depleted intestines. **A-B)** Fly-FUCCI expression in ISCs and EBs. Colored arrows point to cells that are in G1 (green, expressing GFP-E2F1), S (red, expressing RFP-CycB), G2 (yellow, expressing both GFP-E2F1 and RFP-CycB), or unclassified (grey). **(A)** Controls have progenitor cells expressing GFP alone (G1), RFP alone (S), or both GFP and RFP (G2). **(B)** Myt1-depleted progenitors show weak reporter expression or expression of GFP alone. Scale bars = 10 µm. **C)** Quantification of progenitors in each cell cycle phase (control n=15; *myt1^RNAi^* n=14). **D)** Cell cycle profiles of EBs expressing *myt1^RNAi^*. Arrows indicate peaks representing cells in G1, G2 or E1 (endoreplicating cells). Control and *myt1^RNAi^* EB populations were comprised of 26% (s.d. +/-5.3) and 43% (s.d. +/-8.9) G1 phase cells respectively, with G2 populations of 35% (s.d. +/-9.9) and 20% (s.d. +/-8.1) respectively, and endoreplicative (E1) populations of 38% (s.d. +/-10.1) and 38% (s.d. +/-6.0) respectively. Five biological FACS replicates were analyzed for *GBE-Su(H)^ts^* and 4 biological FACS replicates for *GBE-Su(H)^ts^>myt1^RNAi^*, with 15 midguts analyzed per replicate. **E)** Quantification of EB number (esg+, Su(H)-GFP+ cells) in a 100 μm x 100 μm area, one frame anterior to the midgut-hindgut transition zone in control (n=12) and *myt1^RNAi^* flies (n=12). (t test, *** p<0.0005). **F)** Midguts from *myt1/+* controls (n=15) and *myt1* mutants (n=12) stained for Delta/Pros and the EC marker Pdm1. Control gut shows Pdm1 localized to large nuclei (ECs). Pdm1 is absent from large nuclei and instead localizes to small progenitor-like cells, including Delta-positive ISCs (inset), in *myt1* mutant midguts. Scale bars = 30 µm.

Next, we assessed cell cycle dynamics specifically in EBs, by analyzing DNA content in FACS-sorted GFP-positive cells. In this experiment, Myt1 depletion resulted in an accumulation of EBs in G1 phase, with a concomitant decrease in G2 phase cells (Figure 3D). No change was apparent in the proportion of endoreplicating cells, however (Figure 3D). Presumably, this reflects the persistence of pre-existing endoreplicated EBs, during 7 days of *myt1^RNAi^* expression. Notably, loss of Myt1 resulted in a broader endoreplicative peak than in controls, indicating that DNA copy number of the endoreplicating EB population was more variable than normal (Figure 3D). These data are consistent with Myt1 depletion affecting G2 phase of the EB cell cycle, influencing the transition from mitotic cycle to endocycle.

Having shown that loss of Myt1 perturbs EB quiescence, we next assessed EB cell fate using an *esg^ts^*-driven reporter line that differentially labels ISCs and EBs (Martin et al., 2018). EB numbers significantly increased after Myt1 depletion, relative to ISCs (Figures 3E, S3A, and S3B). To quantify the differentiated cell types, we used the *esg^ts^-*driven *Flp-Out* line (hereafter *esgF/O*) to mark progenitor cells and their clones (Jiang et al., 2009). Immunolabeling against Prospero (EE-specific marker) revealed no significant difference in EE numbers between controls and flies expressing *myt1^RNAi^*, at 7 days after clone induction (Figure S3C). However, the proportion of ECs, classified as large cells with polyploid nuclei, decreased in Myt1-depleted clones (Figure S3D). This suggests that progenitors accumulated without differentiating into ECs. We then analyzed *myt1* mutant intestines labeled with the EC marker Pdm1 (Mathur et al., 2010). Strikingly, *myt1* mutants displayed minimal Pdm1 labeling of cells with large nuclei (putative ECs), with strongest labeling instead observed in smaller, progenitor-like cells, including Delta-positive ISCs (Figure 3F). Since Pdm1 is transiently upregulated in mitotic progenitors (Tang et al., 2018), these results are consistent with the hyper-proliferation observed in *myt1* mutants. Moreover, the absence of Pdm1 staining in large cells argues that EB to EC differentiation is perturbed. EB-specific Myt1 depletion therefore disrupts G2 phase arrest and EC specification, causing ectopic proliferation and accumulation of aberrant EBs.

### Myt1 promotes EB mitotic cell cycle exit by inhibiting Cyclin A/Cdk1

EB to EC differentiation is a Notch-dependent process accompanied by transition from mitotic-to-endocycle (Zielke et al., 2014). Cyclin A inhibition was recently shown to be important for transcriptional control of the mitotic-to-endocycle switch (Rotelli et al., 2019). Since Myt1 is a known regulator of Cyclin A/Cdk1 during *Drosophila* male meiosis (Varadarajan et al., 2016), we hypothesized it might act through a similar mechanism during EB differentiation. To test this, we initially examined whether EB mitotic exit was dependent upon inhibition of Cdk1. We compared EB-specific expression of transgenic VFP-labeled reporters for functional Cdk1(WT) with a non-inhibitable mutant form, Cdk1(AF) that drives G2/M checkpoint-arrested cells into mitosis (Ayeni et al., 2014). Mitotic cell numbers increased 5-fold in response to EB-specific expression of Cdk1(AF), relative to functional Cdk1 (Figure 4A). Disruption of Cdk1 inhibitory phosphorylation therefore stimulates aberrant mitotic activity in EBs, consistent with Myt1 inhibition of Cdk1 being required for normal cell cycle exit.

**Figure 4.**
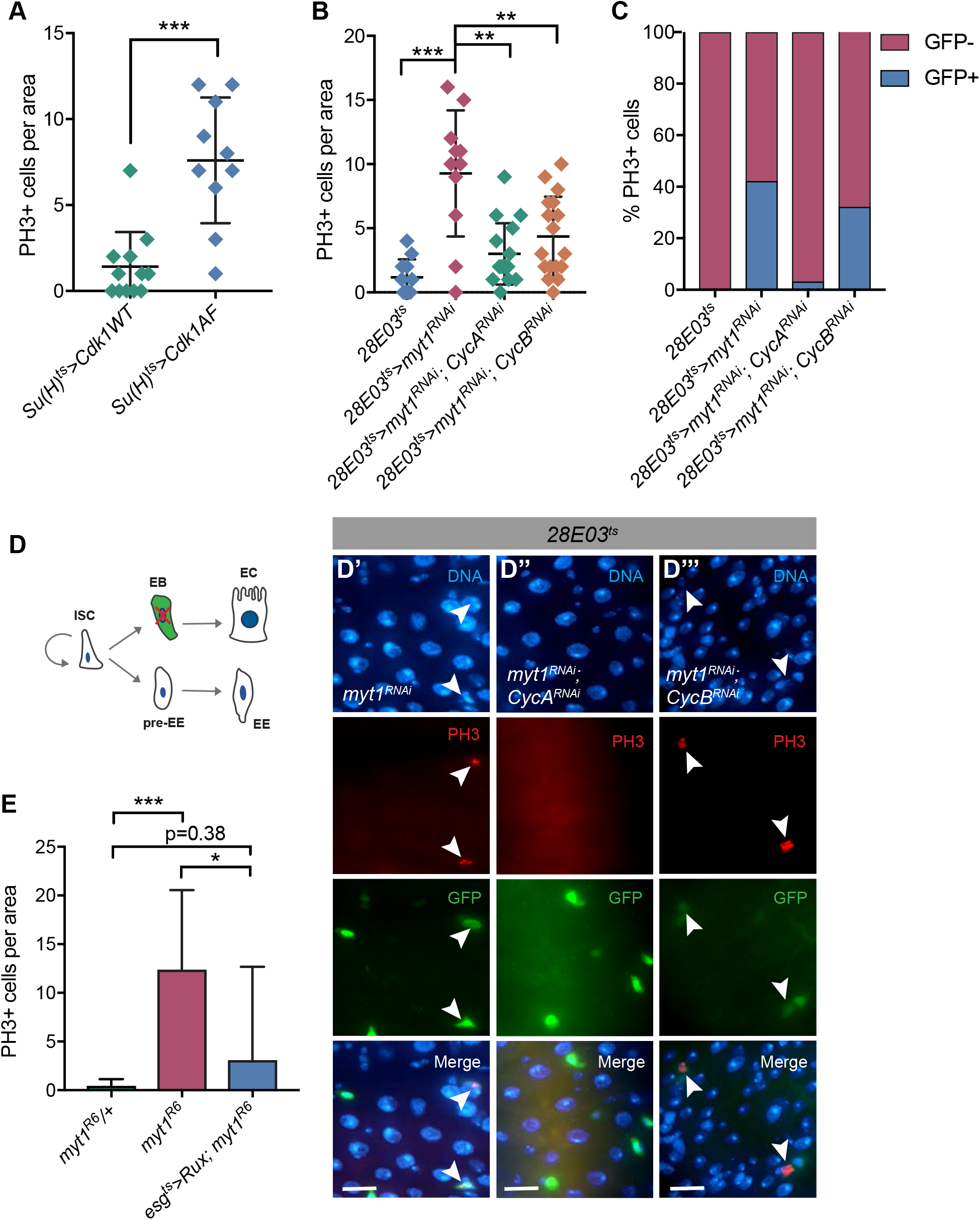
EB entry into mitosis is Cyclin A dependent. **A)** Graph of PH3-positive cells (mitoses) per area in midguts with EBs expressing Cdk1(WT) (n=12) or Cdk1(AF) (n=10). (t test, *** p<0.0005). **B-C)** PH3-labeling was analyzed in the following genotypes: *28E03^ts^* controls (n=11), *28E03^ts^*-driven *myt1^RNAi^* (n=11), or co-expression of *myt1^RNAi^* with *CycA^RNAi^* (n=16), or with *CycB^RNAi^* (n=17). **(B)** Total number of PH3-labeled cells in the gut. **(C)** Proportion of PH3-positive cells present in GFP-positive EBs. **D)** Visualization of GFP-positive EBs expressing *myt1^RNAi^* **(D’)**, *myt1^RNAi^* and *CycA^RNAi^* **(D”)** or *myt1^RNAi^* and *CycB^RNAi^* **(D’”)**. Scale bar = 20 µm. **E)** *esg^ts^*-driven expression of Rux in a *myt1* mutant background rescues the hyper-proliferation phenotype (n=11 for each genotype). Error bars represent +/-SD. (t test, *** p<0.0005; * p<0.05).

We then examined requirements for Cyclin A in the *Drosophila* intestine. Stem cell-specific Cyclin A depletion completely eliminated ISC mitoses, resulting in large, apparently polyploid ISCs, regardless of whether Myt1 was present or not (Figures S4B and S4C). Next, we analyzed whether ectopic EB proliferation in Myt1-depleted guts was Cyclin A-dependent. Flies carrying both *GBE-Su(H)^ts^* and *CycA^RNAi^* transgenes were pupal lethal, so we employed a different EB-specific driver line (*28E03-GAL4*) for this experiment (Lucchetta and Ohlstein, 2017). After making a temperature sensitive, GFP-expressing version of this driver, we compared *myt1^RNAi^* alone to co-expression with either *CycA^RNAi^* or *CycB^RNAi^*. As expected, co-expression of *myt1^RNAi^* with either *CycA^RNAi^* or *CycB^RNAi^* significantly reduced the total number of PH3-positive cells in the gut, compared with expression of *myt1^RNAi^* alone (Figure 4B). Importantly, however, co-depletion of Cyclin A nearly eliminated mitotic EBs relative to Myt1 depletion alone, an effect not seen with co-depletion of Cyclin B (Figures 4C and 4D). We therefore conclude that ectopic mitotic activity in Myt1-depleted EBs reflects a Cyclin A/Cdk1-dependent process. To further test that Myt1 was specifically required to inhibit Cyclin A/Cdk1 we expressed a Cyclin A-specific inhibitor called Roughex (Rux) in a *myt1* mutant background (Foley et al., 1999; Thomas et al., 1994). As predicted, Rux expression also suppressed ectopic proliferation caused by loss of Myt1 activity (Figure 4E). Myt1 inhibition of Cyclin A/Cdk1 is therefore required to prevent ectopic EB mitosis, promoting exit from the mitotic cell cycle and EB to EC differentiation.

## DISCUSSION

In this study we have shown that Myt1 functions as an intrinsic regulator of cell division in *Drosophila*. There are many more mitotic cells in *myt1* mutants and clones grow larger than wild-type controls. Moreover, normally quiescent EBs become mitotic in the absence of Myt1, an anomalous event under homeostatic conditions, but not unprecedented. For example, EBs divide in response to bacterial infection, acting as a rudimentary form of transit-amplifying cell to promote tissue regeneration (Kohlmaier et al., 2015). Additionally, endocyling EBs may undergo amitosis to generate ISCs under starvation conditions (Lucchetta and Ohlstein, 2017). Our observations are therefore consistent with a growing body of evidence that EBs are more adaptable than formerly believed, with mechanisms accommodating EB division or reversion to a stem-like state when the progenitor pool needs replenishment. Since Myt1 loss promotes EB mitosis and progenitor amplification, we speculate that it may be a direct target of differentiation and/or mitogenic cues. If Myt1 activity normally inhibits Cyclin A/Cdk1 to promote EB differentiation, then inhibition of Myt1 would provide a mechanism for enabling Cdk1 to rapidly initiate EB division when tissue regeneration is required.

Canonical EB differentiation involves a mitotic-endocycle (ME) transition. Although polyploid EC-like cells were observed in *myt1* mutants, EB differentiation was clearly disrupted as shown by variations in ploidy levels measured by EB-specific FACS analysis, by the lack of Pdm1-positive ECs and by the lower proportion of ECs observed. In other tissues the ME transition is facilitated by Notch-dependent downregulation of Cdc25 phosphatase and upregulation of Fzr, an APC/C activator (Schaeffer et al., 2004; Shcherbata et al., 2004; Von Stetina et al., 2018). It is therefore plausible that Notch signaling promotes Myt1 repression of Cdk1. Genetic interactions previously implicated Myt1 as a likely downstream target of Notch in the larval eye disc, supporting this idea (Price et al., 2002). Since several other cell types that undergo the ME switch also exhibit ectopic mitosis in *myt1* mutants (Jin et al., 2005; Jin et al., 2008), our findings indicate that Myt1 serves a conserved developmental role promoting the ME switch.

### Cyclin A, G2 phase and cell differentiation

Cyclin A has an established role in promoting mitosis (Vigneron et al., 2018) and has also been implicated in regulation of S phase (Coverley et al., 2002; Rape and Kirschner, 2004) and endoreplication (Sallé et al., 2012). Cell differentiation, however, normally coincides with degradation of mitotic cyclins by the APC/C (Buttitta et al., 2010), with Cyclin A repression acting as a key determinant of the ME switch (Rotelli et al., 2019). EBs are normally in a state of low APC/C activity, with roughly half of these cells in G2 phase (Kohlmaier et al., 2015). How these cells then make the ME transition is unclear. Since EB-specific depletion of Myt1 propels these normally G2 phase-arrested cells into mitosis, we propose that Myt1 inhibition of Cyclin A/Cdk1 promotes the ME switch in the absence of substantial APC/C activity. Cyclin A is also a known inhibitor of Fzr, a key activator of the APC/C (Dienemann and Sprenger, 2004; Reber et al., 2006; Sigrist and Lehner, 1997). Since Fzr was recently shown to mediate the ME switch in the *Drosophila* hindgut during injury repair (Cohen et al., 2018), it would not be surprising if Cyclin A/Cdk1 is an upstream regulator of Fzr in EBs.

Since *myt1* mutant ECs fail to display the mature EC marker Pdm1, our results suggest that Myt1-mediated G2 phase arrest is a critical stage for EBs to exit the mitotic cell cycle and receive differentiation cues. Consistent with this idea, it was recently demonstrated that EB exit from the mitotic cell cycle occurs prior to EC fate commitment (Martin et al., 2018). Moreover, Notch-dependent cell fate decisions are also determined during G2 phase arrest in *Drosophila* sensory organ precursors (Ayeni et al., 2016; Hunter et al., 2016). This suggests that G2 phase is an important and conserved stage for cells to receive differentiation cues. Besides allowing time for differentiation signals to reach neighbouring cells, G2 phase arrest may also provide sufficient time for chromatin remodeling processes associated with the EB to EC transition (García Del Arco et al., 2018). Indeed, alterations in this chromatin remodeling process could conceivably underlie chromosomal abnormalities observed in mitotic *myt1* mutant cells.

In summary, we have identified the metazoan Cdk1 inhibitor Myt1 as a key regulator of mitotic cell cycle exit in EBs and highlight its functional importance as a regulator of EB cell fate decisions under homeostatic conditions. Furthermore, we have shown that Myt1 promotes the ME switch and EB differentiation by inhibition of Cyclin A/Cdk1. Future work should focus on better understanding Cyclin A-mediated processes that must be inhibited by Myt1 to coordinate cell cycle arrest with cell fate differentiation in complex tissues.

## EXPERIMENTAL PROCEDURES

### *Drosophila* Stocks and Genetics

Fly stocks were maintained at 18°C or 25°C on standard cornmeal food. All experiments were performed on mated females at 7 days old (mutants) or after 7 days of transgene expression. For cell type specific expression of transgenes using GAL80^ts^, flies were raised at 18°C and transferred to 29°C 3-5 days post-eclosion. The UAS-GAL4 system was employed for transgene expression in the *Drosophila* intestine. List of GAL4 transcriptional activator driver lines and the cell type(s) in which they induce transgene expression (references in text): *esg^ts^ –* ISC and EB; *esg^ts^; Su(H)-GAL80* – ISC; *GBE^ts^* – EB; *28E03-Gal4* – EB; *Pros^ts^* – EE; *Myo1A^ts^* – EC; *How^ts^* – VM.

### Mosaic analysis with a repressible cell marker (MARCM)

The MARCM technique was used to generate clones of *myt1* mutant cells in an otherwise heterozygous background (Wu and Luo, 2006). Flies were maintained at RT for 4 days post-eclosion, then heat shocked in a 38.5°C water bath for 2 x 30 min separated by 5 min on ice to increase the number of recombination events (fewer events were observed without ice treatment). Flies were left at RT for 7 days post-heat shock before analysis. Clones were defined as adjacent GFP positive cells.

### Immunostaining

Female flies of desired genotype and age were anaesthetized, scored, and put on ice. Flies were transferred to PBS and the intestines were teased out with forceps then transferred to fresh PBS on ice. After dissection, the tissue was fixed in 8% formaldehyde for 25 min. Guts were then rinsed in PBS + 0.2% Triton X-100 (PBT) and incubated in blocking solution of PBT + 3% BSA (PBTB) for 1 hour. Samples were then incubated overnight with 1° antisera diluted in blocking solution. The next day, guts were washed in PBT then incubated with 2° antibody in PBTB. Finally, guts were washed and counterstained with Hoechst 33258 (1:1000 dilution) then straightened out before mounting to control for variation in intestinal folding.

### Microscopy and Image Processing

Z-stack images (0.3-0.5 µm sections) were acquired using a Zeiss Axioskop wide-field microscope equipped with a Retiga Exi camera, or an Olympus IX-81 spinning disc confocal equipped with a Hamamatsu EMCCD (C9100-13) camera. Z-stack images were merged in Volocity 4, exported as TIFFs, then processed with Adobe Photoshop software. All experimental and control images were captured with identical camera settings and identically manipulated in Photoshop (brightness, contrast, and false color manipulations). Quantifications of cell numbers were conducted by manual counting within indicated regions of the gut. Experiments stating the ‘gut’ was analyzed refer to the entire midgut. Where experiments indicate analysis per area, this refers to all of the gut visible in the microscope field, looking just anterior of the midgut-hindgut transition zone (region R4c of the midgut). Analysis of PH3 positive cells was performed in GraphPad Prism using an unpaired t-test with Welch’s correction to control for differences in variance.

### FACS analysis

Guts were dissected and dissociated as in (Dutta et al., 2015). Briefly, guts were incubated in PBS with elastase (1 mg/mL) for 45 min at 27°C, with intermittent pipetting to agitate the tissue. The dissociated guts were then spun down, re-suspended, and filtered through a 20 µm cell strainer (Pluriselect), then analyzed on an Attune Nxt. Propidium iodide was used to select against dead cells, DRAQ5 was used to measure DNA content, and GFP was used to positively select EBs (*GBE-Su(H)^ts^* drives GFP expression specifically in EBs). DNA profiles were produced using FlowJo software.

## RESOURCES

| REAGENT or RESOURCE | SOURCE | IDENTIFIER |
| --- | --- | --- |
| Antibodies | (Dilution) |  |
| Rabbit polyclonal anti-phospho-Histone H3 (Ser10) | Millipore (1:2000) | Cat# 06-570, RRID: AB_310177 |
| Chicken polyclonal anti-GFP | ThermoFisher Scientific (1:2000) | Cat# PA-19533, RRID: AB_1074893 |
| Mouse monoclonal anti-Prospero | DSHB (1:100) | Cat# MR1A, RRID: AB_528440 |
| Mouse monoclonal anti-Delta | DSHB (1:100) | Cat# C594.9B, RRID: AB_528194 |
| Rabbit polyclonal anti-Pdm1 | Yu Cai (1:500) | N/A |
| Goat anti-rabbit IgG (H+L) secondary antibody, Alexa Fluor 568 | ThermoFisher Scientific (1:2000) | Cat# A11011, RRID: AB_143157 |
| Goat anti-mouse IgG (H+L) secondary antibody, Alexa Fluor 568 | ThermoFisher Scientific (1:2000) | Cat# A11004, RRID: AB_2534072 |
| Goat anti-mouse IgG (H+L) secondary antibody, Alexa Fluor 488 | ThermoFisher Scientific (1:2000) | Cat # A32723, RRID: AB_2633275 |
| Goat anti-Chicken IgY (H+L) secondary antibody, Alexa Fluor 488 | ThermoFisher Scientific (1:2000) | Cat# A11039, RRID: AB_142924 |
| Fly Stocks |  |  |
| *myt1<sup>R6</sup>* | Shelagh Campbell | Submitted to BDSC |
| *wee1<sup>ES1</sup>* | Shelagh Campbell | RRID: BDSC_5833 |
| *MARCM80B* | Bruce Edgar | N/A |
| *FRT80B* | Bloomington Drosophila Stock Center | RRID: BDSC_1988 |
| *FRT80B, myt1<sup>R6</sup>* | This study | N/A |
| *esg<sup>ts</sup>* | Edan Foley | N/A |
| *esg-GAL4, UAS-his::CFP, GBE-Su(H)-GFP::nls; tub-GAL80<sup>ts</sup>* | Lucy O'Brien | N/A |
| *esg<sup>ts</sup>; GBE-Su(H)-GAL80* | Bruce Edgar | N/A |
| *esg<sup>ts</sup>F/O* | Bruce Edgar | N/A |
| *GBE-Su(H)<sup>ts</sup>* | Edan Foley | N/A |
| *28E03* | Bloomington Drosophila Stock Center | RRID: BDSC_45546 |
| *Pros-Gal4<sup>ts</sup>* | Bruce Edgar | N/A |
| *Myo1A-GAL4<sup>ts</sup>* | Edan Foley | N/A |
| *How-GAL4<sup>ts</sup>* | Bruce Edgar | N/A |
| *UAS-myt1<sup>RNai</sup>* | VDRC | RRID: FlyBase_FBst0476985 |
| *UAS-EGFP-Myt1* | Shelagh Campbell | RRID: BDSC_65393 |
| *UAS-FUCCI* | BDSC | RRID: BDSC_55122 |
| *UAS-PCNA::GFP* | N/A | N/A |
| *Su(H)-nls-GFP* | Lucie O'Brien | N/A |
| *NRE-GFP* | Benjamin Ohlstein | N/A |
| *UAS-CycA<sup>RNAi</sup>* | Bloomington Drosophila<br>Stock Center | RRID: BDSC_35694 |
| *UAS-CycB<sup>RNAi</sup>* | Bloomington Drosophila<br>Stock Center | RRID: BDSC_34544 |
| *UAS-Rux* | Bloomington Drosophila<br>Stock Center | RRID: BDSC_9166 |

## AUTHOR CONTRIBUTIONS

R.J.W. and S.D.C conceived and designed the experiments. R.J.W and Z.J. performed the experiments. R.J.W and S.D.C. wrote the manuscript.

## ACKNOWLEDGEMENTS

We are grateful to Bruce Edgar, Edan Foley, and Martin Srayko for critically reading this manuscript. We also thank Bruce Edgar, Edan Foley, Lucy O’Brien, and Benjamin Ohlstein for sharing fly lines and reagents, the Flow Cytometry Core at the University of Alberta for help with the FACS experiment, the Bloomington Drosophila Stock Center and Vienna Drosophila Resource Center for fly lines, and the Developmental Studies Hybridoma Bank for antibodies. A Discovery Grant from the Natural Sciences and Engineering Research Council of Canada (SDC), and an NSERC PGS-M fellowship award (RJW) provided funding for this research.

---

**Figure S1. (supports Figure 1).**
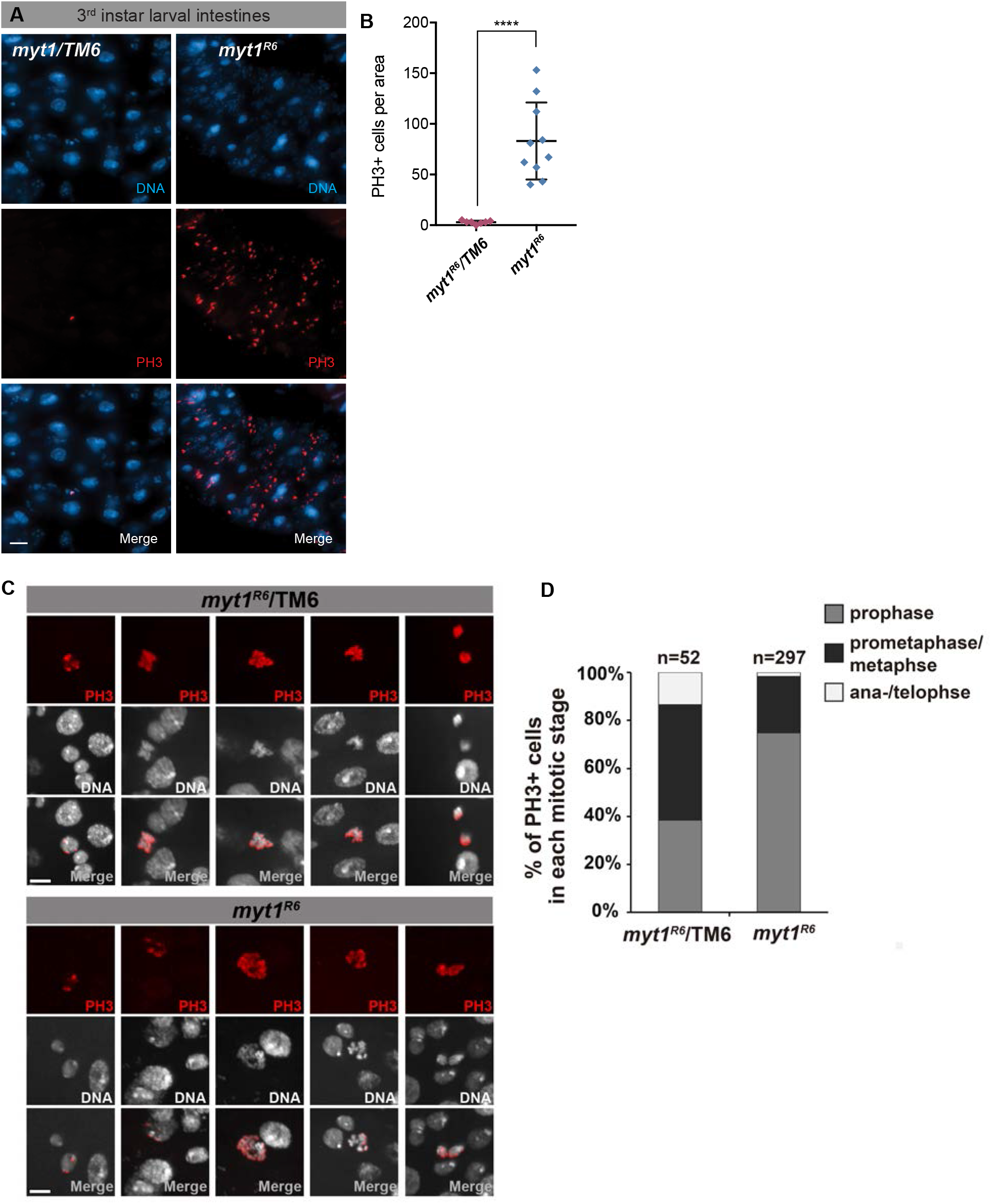
Hyper-proliferation and mitotic analysis of *myt1* mutant intestines. **A)** Representative regions of intestines that were heterozygous or homozygous for a *myt1* null allele, respectively, labeled for PH3. Scale bar = 20 µm. **B)** Quantification of mitoses per frame in control heterozygotes (n=7) and *myt1* mutant (n=10) third instar larval intestines. (t test, **** p<0.00005). Error bars represent +/-SD. **C)** Examples of mitotic stages observed in *myt1* heterozygous and mutant intestines. **D)** Quantification of mitotic stages within *myt1/+* and *myt1* mutant intestines, determined by the morphology of PH3 positive nuclei.

**Figure S2. (supports Figure 2).**
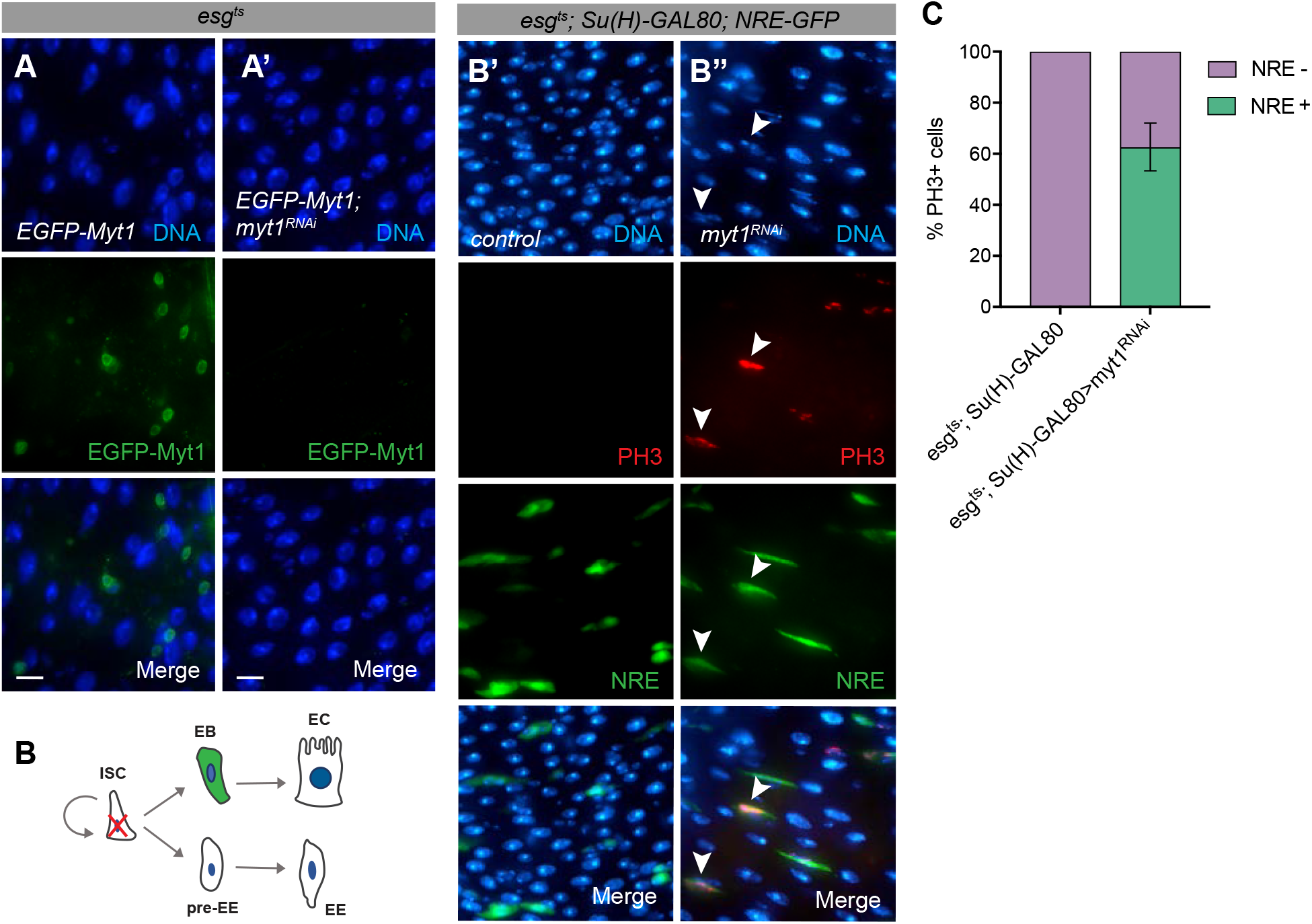
Membrane-localized Myt1 prevents mitosis in EBs. **A)** *esg^ts^*-driven EGFP-Myt1 expressed in progenitor cells localizes to the endomembrane/nuclear envelope, whereas no GFP signal is detected upon co-expression of EGFP-Myt1 and *myt1^RNAi^* **(A’)**. Guts were analyzed 7 days after temperature shift. Scale bar = 10 µm. **B-C)** RNAi against *myt1* is expressed in ISCs and EBs are marked by an NRE-GFP reporter. **(B”)** PH3-positive EBs are not observed in control guts **(B’)** but were seen in guts with Myt1-depleted ISCs **(B’)**. Scale bar = 10 µm. **(C)** Quantification of mitotic cells that were also positive for the EB marker NRE-GFP.

**Figure S3. (supports Figure 3).**
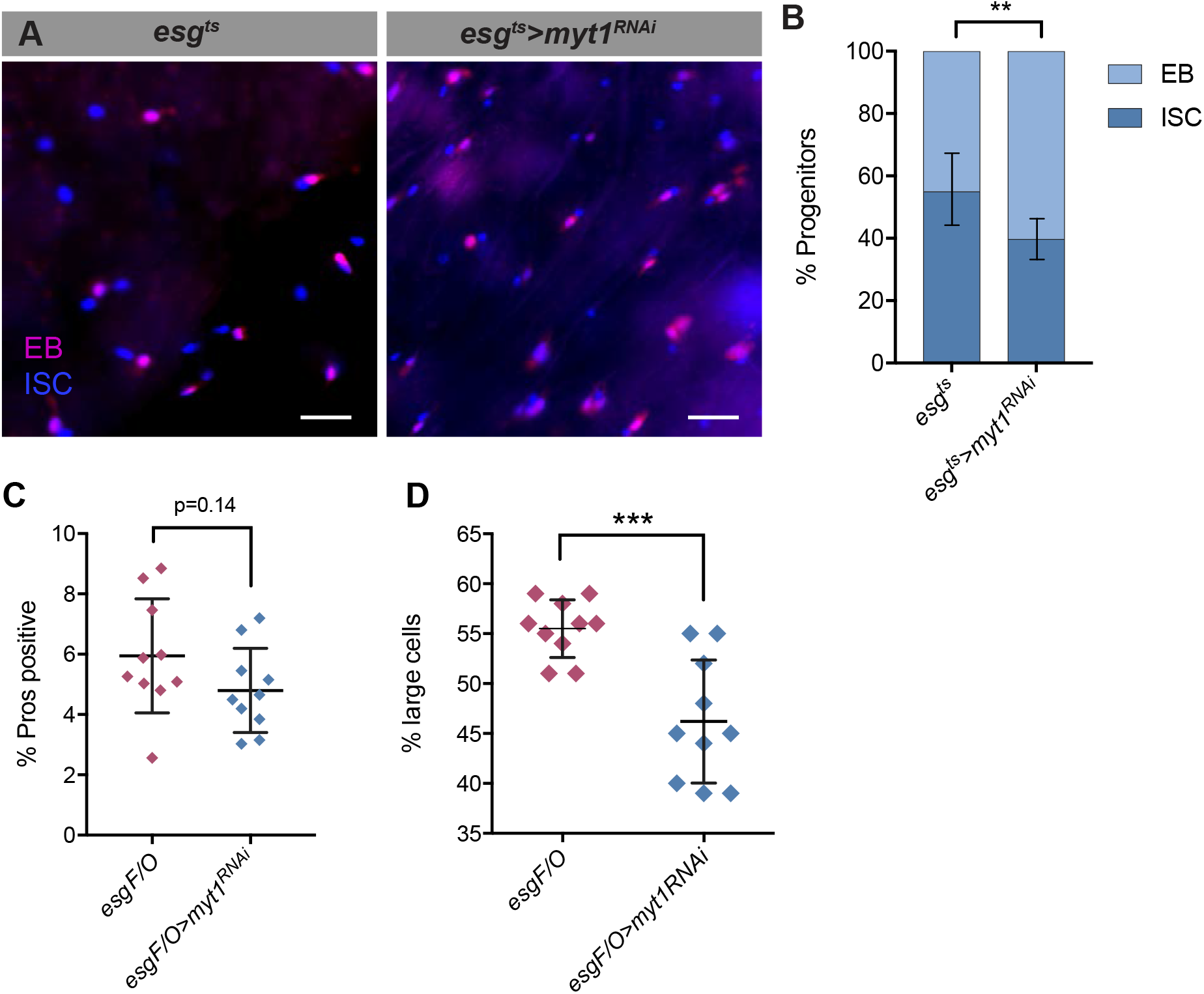
Myt1 loss affects EB to EC differentiation. **A)** Visualization of ISCs and EBs using *esgGAL4, UAS-Histone::CFP*, *GBE-Su(H)-GFP:nls; tub-GAL80^ts^* in control and Myt1 knockdown flies, where ISCs are CFP-positive (blue) and EBs are CFP and GFP-positive (magenta). Scale bars = 15 µm. **B)** Quantification of relative ISC and EB numbers, from (A). **C)** The percentage of EEs (Pros-positive nuclei) in GFP-positive clones does not change in *esgF/O>myt1^RNAi^* flies (n=10), relative to controls (n=10). **D)** The percentage of ECs (large nuclei) in GFP-positive clones decreases upon Myt1 depletion. (t test, *** p<0.0005). Error bars represent +/-SD.

**Figure S4. (supports Figure 4).**
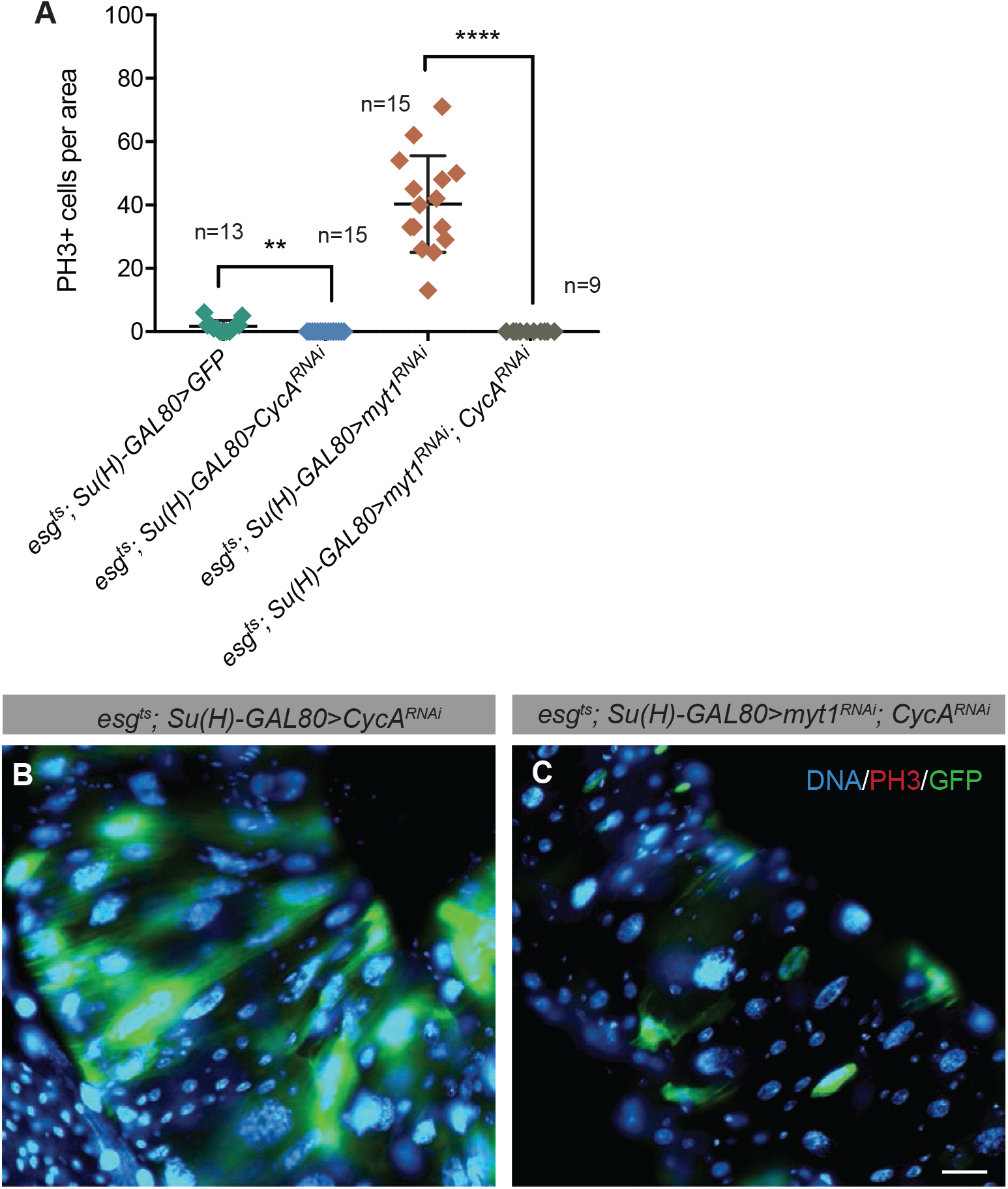
Cyclin A is essential for mitosis in the *Drosophila* intestine. **A)** Quantification of PH3-positive cells in posterior midguts at 7 days after ISC-specific transgene expression. (t test, ** p<0.005; **** p<0.00005) **B)** ISC-specific Cyclin A depletion produces large polyploid cells (stem cells are GFP-positive), with no observable mitoses. **C)** Co-expression of *myt1^RNAi^* and *CycA^RNAi^* also results in large polyploid cells with no observable mitoses. Scale bar = 32 μm.

## REFERENCES

Ayeni, J.O., Audibert, A., Fichelson, P., Srayko, M., Gho, M., and Campbell, S.D. (2016) G2 phase arrest prevents bristle progenitor self-renewal and synchronizes cell division with cell fate differentiation. Development, 143, 1160–1169.

Ayeni, J.O., Varadarajan, R., Mukherjee, O., Stuart, D.T., Sprenger, F., Srayko, M., and Campbell, S.D. (2014) Dual phosphorylation of cdk1 coordinates cell proliferation with key developmental processes in Drosophila. Genetics, 196, 197–210.

Buchon, N., Broderick, N.A., Poidevin, M., Pradervand, S., and Lemaitre, B. (2009) Drosophila intestinal response to bacterial infection: activation of host defense and stem cell proliferation. Cell Host Microbe, 5, 200–211.

Buttitta, L.A., and Edgar, B.A. (2007) Mechanisms controlling cell cycle exit upon terminal differentiation. Curr Opin Cell Biol, 19, 697–704.

Buttitta, L.A., Katzaroff, A.J., and Edgar, B.A. (2010) A robust cell cycle control mechanism limits E2F-induced proliferation of terminally differentiated cells in vivo. J Cell Biol, 189, 981–996.

Cohen, E., Allen, S.R., Sawyer, J.K., and Fox, D.T. (2018) Fizzy-related dictates a cell cycle switch during organ repair and tissue growth responses in the Drosophila hindgut. Elife, 7,

Coverley, D., Laman, H., and Laskey, R.A. (2002) Distinct roles for cyclins E and A during DNA replication complex assembly and activation. Nat Cell Biol, 4, 523–528.

Dienemann, A., and Sprenger, F. (2004) Requirements of cyclin a for mitosis are independent of its subcellular localization. Curr Biol, 14, 1117–1123.

Dutta, D., Buchon, N., Xiang, J., and Edgar, B.A. (2015) Regional Cell Specific RNA Expression Profiling of FACS Isolated Drosophila Intestinal Cell Populations. Curr Protoc Stem Cell Biol, 34, 2F.2.1–14.

Fasulo, B., Koyama, C., Yu, K.R., Homola, E.M., Hsieh, T.S., Campbell, S.D., and Sullivan, W. (2012) Chk1 and Wee1 kinases coordinate DNA replication, chromosome condensation, and anaphase entry. Mol Biol Cell, 23, 1047–1057.

Foley, E., O’Farrell, P.H., and Sprenger, F. (1999) Rux is a cyclin-dependent kinase inhibitor (CKI) specific for mitotic cyclin-Cdk complexes. Curr Biol, 9, 1392–1402.

García Del Arco, A., Edgar, B.A., and Erhardt, S. (2018) In Vivo Analysis of Centromeric Proteins Reveals a Stem Cell-Specific Asymmetry and an Essential Role in Differentiated, Non-proliferating Cells. Cell Rep, 22, 1982–1993.

Hendzel, M.J., Wei, Y., Mancini, M.A., Van Hooser, A., Ranalli, T., Brinkley, B.R., Bazett-Jones, D.P., and Allis, C.D. (1997) Mitosis-specific phosphorylation of histone H3 initiates primarily within pericentromeric heterochromatin during G2 and spreads in an ordered fashion coincident with mitotic chromosome condensation. Chromosoma, 106, 348–360.

Hunter, G.L., Hadjivasiliou, Z., Bonin, H., He, L., Perrimon, N., Charras, G., and Baum, B. (2016) Coordinated control of Notch/Delta signalling and cell cycle progression drives lateral inhibition-mediated tissue patterning. Development, 143, 2305–2310.

Jiang, H., Patel, P.H., Kohlmaier, A., Grenley, M.O., McEwen, D.G., and Edgar, B.A. (2009) Cytokine/Jak/Stat signaling mediates regeneration and homeostasis in the Drosophila midgut. Cell, 137, 1343–1355.

Jin, Z., Homola, E., Tiong, S., and Campbell, S.D. (2008) Drosophila myt1 is the major cdk1 inhibitory kinase for wing imaginal disc development. Genetics, 180, 2123–2133.

Jin, Z., Homola, E.M., Goldbach, P., Choi, Y., Brill, J.A., and Campbell, S.D. (2005) Drosophila Myt1 is a Cdk1 inhibitory kinase that regulates multiple aspects of cell cycle behavior during gametogenesis. Development, 132, 4075–4085.

Kohlmaier, A., Fassnacht, C., Jin, Y., Reuter, H., Begum, J., Dutta, D., and Edgar, B.A. (2015) Src kinase function controls progenitor cell pools during regeneration and tumor onset in the Drosophila intestine. Oncogene, 34, 2371–2384.

Lee, T., and Luo, L. (1999) Mosaic analysis with a repressible cell marker for studies of gene function in neuronal morphogenesis. Neuron, 22, 451–461.

Lee, W.C., Beebe, K., Sudmeier, L., and Micchelli, C.A. (2009) Adenomatous polyposis coli regulates Drosophila intestinal stem cell proliferation. Development, 136, 2255–2264.

Lehner, C.F., and O’Farrell, P.H. (1990) The roles of Drosophila cyclins A and B in mitotic control. Cell, 61, 535–547.

Lin, G., Xu, N., and Xi, R. (2008) Paracrine Wingless signalling controls self-renewal of Drosophila intestinal stem cells. Nature, 455, 1119–1123.

Lucchetta, E.M., and Ohlstein, B. (2017) Amitosis of Polyploid Cells Regenerates Functional Stem Cells in the Drosophila Intestine. Cell Stem Cell, 20, 609–620.e6.

Mathur, D., Bost, A., Driver, I., and Ohlstein, B. (2010) A transient niche regulates the specification of Drosophila intestinal stem cells. Science, 327, 210–213.

Martin, J.L., Sanders, E.N., Moreno-Roman, P., Jaramillo Koyama, L.A., Balachandra, S., Du, X., and O’Brien, L.E. (2018) Long-term live imaging of the Drosophila adult midgut reveals real-time dynamics of division, differentiation and loss. Elife, 7,

Micchelli, C.A., and Perrimon, N. (2006) Evidence that stem cells reside in the adult Drosophila midgut epithelium. Nature, 439, 475–479.

Ohlstein, B., and Spradling, A. (2006) The adult Drosophila posterior midgut is maintained by pluripotent stem cells. Nature, 439, 470–474.

Ohlstein, B., and Spradling, A. (2007) Multipotent Drosophila intestinal stem cells specify daughter cell fates by differential notch signaling. Science, 315, 988–992.

Pimentel, A.C., and Venkatesh, T.R. (2005) rap gene encodes Fizzy-related protein (Fzr) and regulates cell proliferation and pattern formation in the developing Drosophila eye-antennal disc. Dev Biol, 285, 436–446.

Price, D., Rabinovitch, S., O’Farrell, P.H., and Campbell, S.D. (2000) Drosophila wee1 has an essential role in the nuclear divisions of early embryogenesis. Genetics, 155, 159–166.

Price, D.M., Jin, Z., Rabinovitch, S., and Campbell, S.D. (2002) Ectopic expression of the Drosophila Cdk1 inhibitory kinases, Wee1 and Myt1, interferes with the second mitotic wave and disrupts pattern formation during eye development. Genetics, 161, 721–731.

Rape, M., and Kirschner, M.W. (2004) Autonomous regulation of the anaphase-promoting complex couples mitosis to S-phase entry. Nature, 432, 588–595.

Reber, A., Lehner, C.F., and Jacobs, H.W. (2006) Terminal mitoses require negative regulation of Fzr/Cdh1 by Cyclin A, preventing premature degradation of mitotic cyclins and String/Cdc25. Development, 133, 3201–3211.

Rotelli, M.D., Policastro, R.A., Bolling, A.M., Killion, A.W., Weinberg, A.J., Dixon, M.J., Zentner, G.E., Walczak, C.E., Lilly, M.A., and Calvi, B.R. (2019) A Cyclin A-Myb-MuvB-Aurora B network regulates the choice between mitotic cycles and polyploid endoreplication cycles. PLoS Genet, 15, e1008253.

Sallé, J., Campbell, S.D., Gho, M., and Audibert, A. (2012) CycA is involved in the control of endoreplication dynamics in the Drosophila bristle lineage. Development, 139, 547–557.

Schaeffer, V., Althauser, C., Shcherbata, H.R., Deng, W.M., and Ruohola-Baker, H. (2004) Notch-dependent Fizzy-related/Hec1/Cdh1 expression is required for the mitotic-to-endocycle transition in Drosophila follicle cells. Curr Biol, 14, 630–636.

Shcherbata, H.R., Althauser, C., Findley, S.D., and Ruohola-Baker, H. (2004) The mitotic-to-endocycle switch in Drosophila follicle cells is executed by Notch-dependent regulation of G1/S, G2/M and M/G1 cell-cycle transitions. Development, 131, 3169–3181.

Sigrist, S.J., and Lehner, C.F. (1997) Drosophila fizzy-related down-regulates mitotic cyclins and is required for cell proliferation arrest and entry into endocycles. Cell, 90, 671–681.

Stumpff, J., Duncan, T., Homola, E., Campbell, S.D., and Su, T.T. (2004) Drosophila Wee1 kinase regulates Cdk1 and mitotic entry during embryogenesis. Curr Biol, 14, 2143–2148.

Tang, X., Zhao, Y., Buchon, N., and Engström, Y. (2018) The POU/Oct Transcription Factor Nubbin Controls the Balance of Intestinal Stem Cell Maintenance and Differentiation by Isoform-Specific Regulation. Stem Cell Reports, 10, 1565–1578.

Thomas, B.J., Gunning, D.A., Cho, J., and Zipursky, L. (1994) Cell cycle progression in the developing Drosophila eye: roughex encodes a novel protein required for the establishment of G1. Cell, 77, 1003–1014.

Varadarajan, R., Ayeni, J., Jin, Z., Homola, E., and Campbell, S.D. (2016) Myt1 inhibition of Cyclin A/Cdk1 is essential for fusome integrity and premeiotic centriole engagement in Drosophila spermatocytes. Mol Biol Cell, 27, 2051–2063.

Vigneron, S., Sundermann, L., Labbé, J.C., Pintard, L., Radulescu, O., Castro, A., and Lorca, T. (2018) Cyclin A-cdk1-Dependent Phosphorylation of Bora Is the Triggering Factor Promoting Mitotic Entry. Dev Cell, 45, 637–650.e7.

Von Stetina, J.R., Frawley, L.E., Unhavaithaya, Y., and Orr-Weaver, T.L. (2018) Variant cell cycles regulated by Notch signaling control cell size and ensure a functional blood-brain barrier. Development, 145,

Wang, L., Zeng, X., Ryoo, H.D., and Jasper, H. (2014) Integration of UPRER and oxidative stress signaling in the control of intestinal stem cell proliferation. PLoS Genet, 10, e1004568.

Wu, J.S., and Luo, L. (2006) A protocol for mosaic analysis with a repressible cell marker (MARCM) in Drosophila. Nat Protoc, 1, 2583–2589.

Zeng, X., Chauhan, C., and Hou, S.X. (2010) Characterization of midgut stem cell- and enteroblast-specific Gal4 lines in drosophila. Genesis, 48, 607–611.

Zielke, N., Korzelius, J., van Straaten, M., Bender, K., Schuhknecht, G.F., Dutta, D., Xiang, J., and Edgar, B.A. (2014) Fly-FUCCI: A versatile tool for studying cell proliferation in complex tissues. Cell Rep, 7, 588–598.

